# Spatial mRNA expression patterns of orexin receptors in the dorsal hippocampus

**DOI:** 10.1101/2024.04.15.589483

**Authors:** Gina Marie Krause, Lara Mariel Chirich Barreira, Anne Albrecht

**Author notes:** ***corresponding author:*** Anne Albrecht, Institute of Anatomy, Otto-von-Guericke-University, Leipziger Str. 44, 39120 Magdeburg, Germany.

## Abstract

Orexins are wake-promoting neuropeptides that originate from hypothalamic neurons projecting to widespread brain areas throughout the central nervous system. They modulate various physiological functions via their orexin 1 (OXR1) and 2 (OXR2) receptors, including sleep-wake rhythm but also cognitive functions such as memory formation.

Here, we provide a detailed analysis of OXR1 and OXR2 mRNA expression profiles in the dorsal hippocampus as a key region for memory formation, using RNAscope® multiplex *in situ* hybridization. Interconnected subareas relevant for cognition and memory such as the medial prefrontal cortex and the nucleus reuniens of the thalamus were assessed as well. Both receptor types display distinct profiles, with the highest percentage of OXR1 mRNA-positive cells in the hilus of the dentate gyrus. Here, the content of OXR1 mRNA per cell was slightly modulated at selected time points over a 12h light/ 12 dark light phase. Using RNAScope® and quantitative polymerase chain reaction approaches, we began to address a cell-type specific expression of OXR1 in hilar GABAergic interneurons.

The distinct expression profiles of both receptor subtypes within hippocampal subareas and circuits provide an interesting basis for future interventional studies on orexin receptor function in spatial and contextual memory.

## 1. IntroducBon

The neuropeptide orexin has been prompted as the “key to a healthy life” ^1^. Indeed, while well described as a wake-promoting circadian regulator in the brain, recent evidence suggests that the orexinergic system is involved in different forms of learning and memory and its dysfunction may contribute to neurodegenerative diseases such as Alzheimer’s ^2^. Thus, understanding the expression and function of orexin and its receptors in the hippocampus has recently gained interest ^3^.

Orexin, also known as hypocretin, is a neuropeptide produced in a group of neurons located mainly in the lateral hypothalamus from a precursor protein called prepro-orexin (131 amino acids) that is enzymatically cleaved into orexin-A (33 amino acids) and orexin-B (28 amino acids) ^4,5^. The functional neuropeptides orexin-A and B are packed into vesicles and transferred to numerous target sites throughout the brain, including the hippocampus and other structures related to learning and memory ^6^. Regional actions of orexin within its target areas are exerted via two receptor subtypes, orexin receptor 1 and 2 (OXR1 and OXR2). While OXR1 binds only orexin-A, OXR2 can bind both orexin isoforms with equal affinity. Both receptor subtypes belong to the subclass of heterotrimeric G-protein-coupled receptors (GPCRs). OXR1 builds a complex with Gq proteins, triggering calcium influx via an inositoltriphosphate (IP3)-dependent signalling cascade, thereby activating the targeted cell. OXR2 can be associated to both Gq and inhibitory Gi/o proteins, the later causing hyperpolarization via potassium efflux ^7^. The activation of orexinergic neurons in the lateral hypothalamus is highest during the active phase and lowest towards the end of the inactive phase (i.e. dark vs. light phase for nocturnal model animals such as mice and rats) and before the transition from NREM to REM sleep phases ^8,9^. Accordingly, intracerebral application of orexin-A in rats induces waking and increases the time awake ^10^, while in orexin knock-out mice fragmented sleep and cataplectic episodes during the active phase are observed ^11^. Similar findings in different transgenic animals and with pharmacological approaches prompted the appreciation of orexin as a wake promoting neuropeptide and orexin knock-out models as valuable animal models to study sleep-related diseases such as narcolepsy ^12^. Moreover, these animal models suggest a prominent role of orexin in learning and memory as well. Orexin knock-out mice display deficits in spatial working memory, spatial object location tasks ^13^ and in contextual and cued fear memory ^14^. Pharmacological studies have helped to link local orexin functions to its receptor subtypes. OXR1 signalling in the Cornu ammonis (CA) 1 and dentate gyrus (DG) subfield of the hippocampus has been linked to spatial learning and memory ^15,16^. A dysregulation in the hippocampal expression of orexin receptors was also observed in mouse models for post-traumatic stress disorder and in Alzheimer’s disease ^17,18^, both are disorders that show features of dysfunctional or impaired hippocampus-dependent memory. Recent evidence suggest that orexin drives the activity of the medial prefrontal cortex (mPFC) ^19,20^. Indeed, the hippocampus and the mPFC are functionally coupled during working memory performance and for the long-term storage, retrieval and organization of spatial and contextual fear memories. Anatomically, direct projections of the ventral hippocampus exist next to bidirectional connections via relay stations, such as the nucleus reuniens (NRe) that reciprocally connects the mPFC with the CA1 and subiculum of the hippocampus ^21^.

Despite the broad utilization of mice in investigating these functions connected to the hippocampus and its interactive circuits, systematic studies of orexin receptor expression profiles were performed mainly in rats ^22–25^ and only recently a comprehensive study assessing OXR1 and OXR2 mRNA throughout the mouse brain using high-sensitivity *in situ* hybridization was performed ^26^. To complement this data, we now set out to investigate the OXR1 and OXR2 mRNA expression profiles in the dorsal hippocampus and in interconnected areas relevant for memory and cognition, such as the PFC and the NRe in greater detail. To this end, we analysed distinct sublayers of the dorsal hippocampal regions DG, CA1 and CA3 as well as subareas of the mPFC. Since orexinergic neurons display a diurnal rhythm of activity, peaking during the waking phase ^8^, we compared the expression of OXR1 and OXR2 at different time points over a 12h dark/ 12h light cycle. We used RNAScope, a commercially available high-sensitive fluorescence in situ hybridisation technique that allows to assess the co-expression of OXR1 and OXR2 mRNA within the same neurons, and applied a semiquantitative analysis of their expression patterns in relation to the overall cell densities in our target regions. Cell-type specific expression profiles were assessed for the hilus of the DG, which contains diverse populations of interneurons regulating spatial and contextual fear memory.

## 2. Results

### 2.1 Layer-specific hippocampal expression profiles of OXR1 and OXR2 mRNA

To investigate whether OXR1 and OXR2 mRNA expression profiles within different layers of the dorsal hippocampus depend on the time of brain preparation in a 12h dark/ 12h light cycle, a mixed-model analysis was performed for the factors Zeitgeber time (ZT) and layer for each subarea. For sublayers of the dorsal DG, CA3 and CA1, only layer-specific expression effects were significant, but not the time point of preparation indicated by ZT nor the interactions of ZT and layer within the subregions (Suppl. Fig. S1 and suppl. table S1).

For a further analysis of expression profiles, the ZT groups were therefore combined and in the next step, a mixed-model analysis was performed for the factors receptor subtype (OXR1 and OXR2) and sublayer in order to investigate their differential expression in DG, CA3 an CA1. An interaction of layer x receptor type and a main effect of receptor type were observed in all three hippocampal subregions (see Table 1). Tukey’s multiple comparisons post hoc tests revealed an increased number of OXR1(+) cells compared to OXR2(+) cells in the hilus of the DG (Fig. 1B; p<0.0001), while more OXR2(+) cells than OXR1(+) cells were observed in the molecular layer (ML) of the DG, however with an overall much lower expression level of both receptors in the ML compared to the hilus. Similarly, higher numbers of OXR2(+) compared to OXR1(+) cells were observed for the *stratum pyramidale* (SP) of CA3 and CA1 as well as for the *stratum radiale* (SR) and the *stratum oriens* (SO) of CA1 (Fig. 1C, D; see Table 1 for details on statistics).

**Fig. 1:**
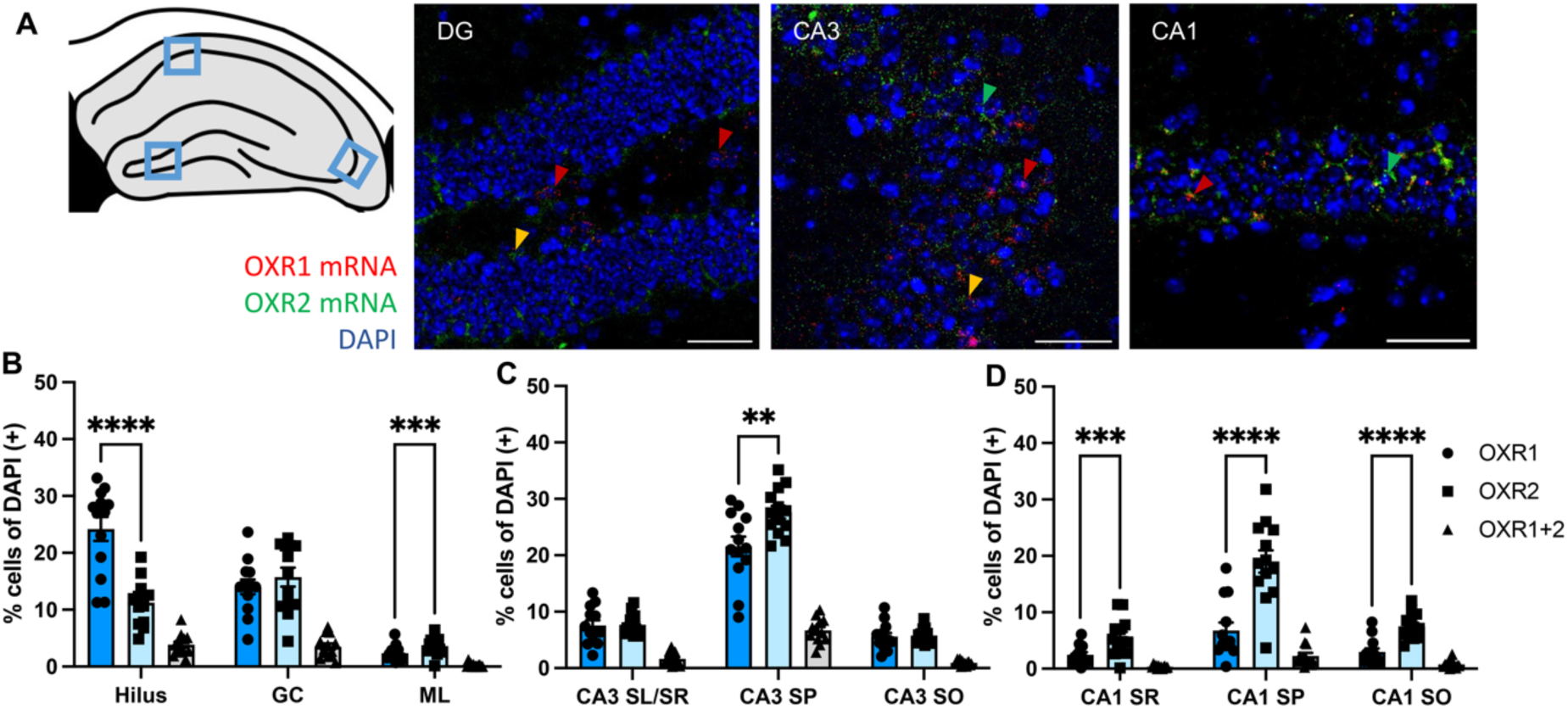
Orexin receptor mRNA expression profiles in hippocampal sublayers. (**A**) Exemplary images of OXR1 and OXR2 mRNA detection in the dorsal hippocampus (scale bar: 50 µM), imaging position indicated in the scheme. Arrowheads indicate cells positive for OXR1 (red), OXR2 (green) or OXR1+OXR2 mRNA (orange). (**B-D**) The expression levels of OXR1 and OXR2 mRNA show layer-specific patterns in the dorsal hippocampus with increased expression of OXR1 vs. OXR2 in the hilus of the dorsal DG. OXR2 expression levels were increased compared to OXR1 in the SP of CA3 and in all sublayers analysed of the CA1. Cells expressing both OXR1 and OXR2 mRNA were barely detected in all sublayers. DG, dentate gyrus; GC, granular cell layer; ML, molecular cell layer; CA, *cornu ammonis*; SO, *stratum oriens*; SP, *stratum pyramidale*; SR, *stratum radiatum*; SL, *stratum lucidum*. Graphs show mean ± SEM of mice pooled over all time points (n=13) since no diurnal differences in OXR1 or OXR2 mRNA expression were evident. ** significant difference between receptors, ** p<0.01; *** p<0.001; **** p<0.0001.

**Table 1:** Statistical results for mixed model analysis in hippocampal sublayers. Tukey’s post hoc comparison for percentage of cells containing OXR1 vs. OXR2 mRNA. DG, dentate gyrus; GC, granular cell layer; ML, molecular cell layer; CA, cornu ammonis; SO, stratum oriens; SP, stratum pyramidale; SR, stratum radiatum; SL, stratum lucidum.

|  |  | Mixed model analysis | Post hoc<br>OXR1 vs. 2 | Post hoc layer<br>OXR1 | Post hoc layer<br>OXR2 |
| --- | --- | --- | --- | --- | --- |
| **DG** | layer x<br>receptor | $F(1.665, 19.98) = 50.22$;<br>$p < 0.0001$ | Hilus: $p < 0.0001$;<br>GC: $p = 0.2289$;<br>ML: $p = 0.0002$ | Hilus vs. GC:<br>$p < 0.0001$;<br>Hilus vs. ML:<br>$p < 0.0001$;<br>GC vs. ML:<br>$p < 0.0001$ | Hilus vs. GC:<br>$p = 0.0276$;<br>Hilus vs. ML:<br>$p < 0.0001$;<br>GC vs. ML:<br>$p < 0.0001$ |
| | layer | $F(1.921, 23.06) = 115.4$;<br>$p < 0.0001$ | | | |
| | receptor | $F(1, 12) = 14.54$;<br>$p = 0.0025$ | | | |
| **CA3** | layer x<br>receptor | $F(1.236, 14.84) = 15.98$;<br>$p = 0.0007$ | CA3 SR/SL:<br>$p = 0.9021$;<br>CA3 SP:<br>$p = 0.0057$;<br>CA3 SO:<br>$p = 0.7706$ | CA3 SL/SR vs. CA3 SP:<br>$p < 0.0001$;<br>CA3 SL/SR vs. CA3<br>SO: $p = 0.0050$;<br>CA3 SP vs. CA3 SO:<br>$p < 0.0001$ | CA3 SL/SR vs. CA3<br>SP: $p < 0.0001$;<br>CA3 SL/SR vs. CA3<br>SO: $p = 0.0005$;<br>CA3 SP vs. CA3 SO:<br>$p < 0.0001$ |
| | layer | $F(1.178, 14.13) = 351.8$;<br>$p < 0.0001$ | | | |
| | receptor | $F(1, 12) = 4.174$;<br>$p = 0.0636$ | | | |
| **CA1** | layer x<br>receptor | $F(1.513, 18.15) = 39.43$;<br>$p < 0.0001$ | CA1 SR:<br>$p = 0.0008$;<br>CA1 SP:<br>$p < 0.0001$;<br>CA1 SO:<br>$p < 0.0001$ | CA1 SR vs. CA1 SP:<br>$p = 0.0041$;<br>CA1 SR vs. CA1 SO:<br>$p = 0.4541$;<br>CA1 SP vs. CA1 SO:<br>$p = 0.0044$ | CA1 SR vs. CA1 SP:<br>$p < 0.0001$;<br>CA1 SR vs. CA1 SO:<br>$p = 0.0146$;<br>CA1 SP vs. CA1 SO:<br>$p < 0.0001$ |
| | layer | $F(1.108, 13.30) = 57.99$;<br>$p < 0.0001$ | | | |
| | receptor | $F(1, 12) = 71.25$;<br>$p < 0.0001$ | | | |

In all three regions, only a small fraction of cells expressed both receptor subtypes (below 10%, see Fig. 1 and Suppl. Fig. S1, Suppl. Table S1 for statistical details).

### 2.2 Subregion-specific expression profiles of OXR1 and OXR2 mRNA for the PFC-Hippocampus network

As an important site for hippocampal interaction in cognition and memory, OXR1 and OXR2 mRNA expression profiles were investigated in mPFC subregions as well as in the NRe, an important PFC-to-hippocampus relay station. Again, no ZT-dependent effects were observed when employing a mixed-model analysis for ZT and subarea (Suppl. Fig. S2 and Suppl. Table S2 for overview).

After combining the ZT groups, the cell counts of OXR1(+) and OXR2(+) neurons were compared for the mPFC by a mixed model analysis for the factors subarea and receptor subtype, revealing an effect of receptor type (see Table 2 for details on statistics). Tukey’s multiple comparisons post hoc tests demonstrated an increased number of OXR2(+) compared to OXR1(+) cells for all mPFC subareas cingulate (Cg), prelimbic (PL) and infralimbic (IL) cortex (Fig. 2B; p<0.0001). Receptor mRNA expression counts in the NRe were compared using a paired T-test, which failed to reach significance (Fig. 2C; p=0.0613), indicating similarly high expression levels for OXR1 and OXR2 in these structures.

**Fig. 2:**
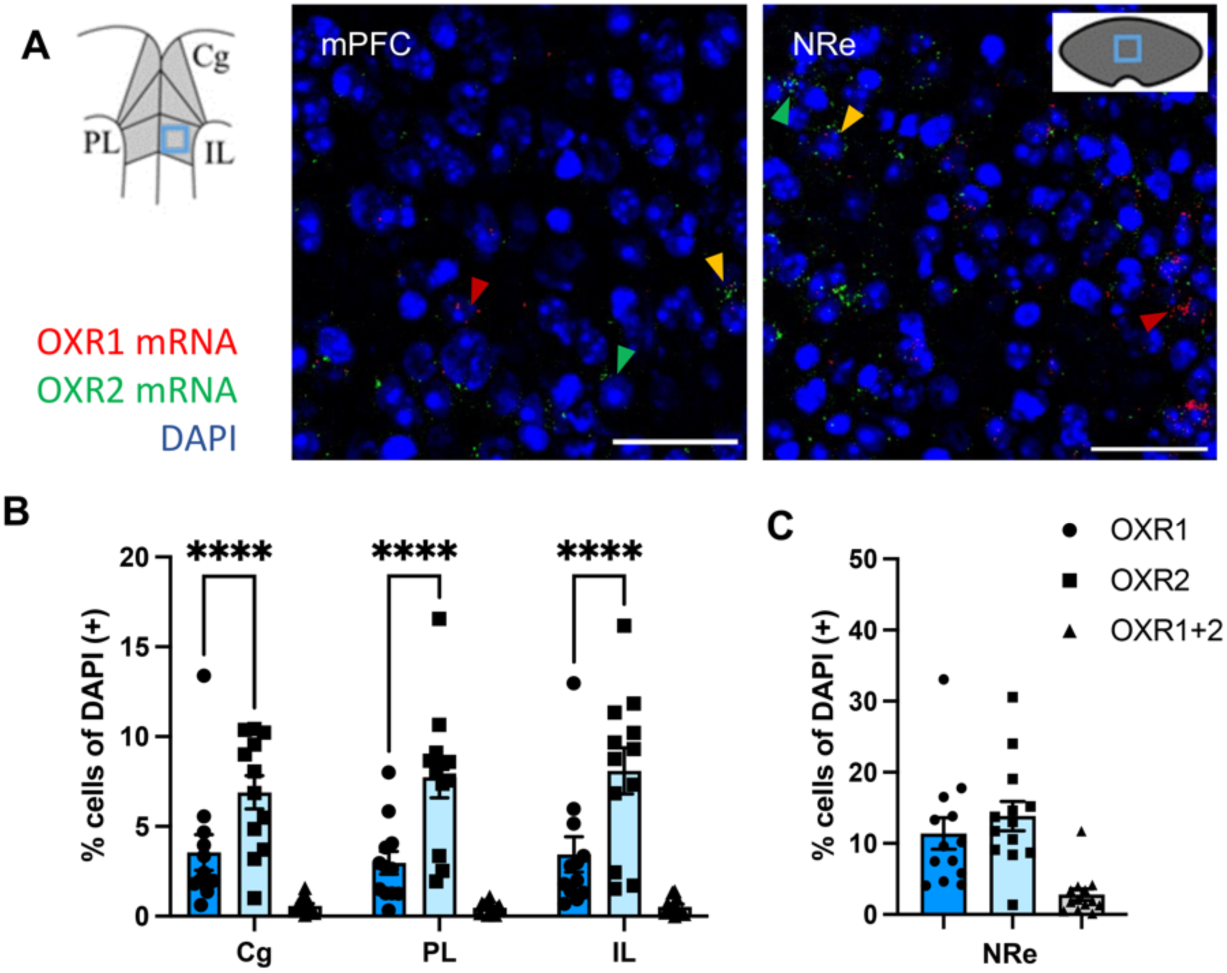
Orexin receptor mRNA expression profiles in the mPFC subregions and the nucleus reuniens relay staLon to the hippocampus. (A) Exemplary images of OXR1 and OXR2 mRNA detection in the mPFC and NRe (scale bar: 50 µM), imaging position indicated in the schemes. Arrowheads indicate cells positive for OXR1 (red), OXR2 (green) or OXR1+OXR2 mRNA (orange). (B) All subregions of the mPFC show higher OXR2 expression density compared to OXR1. (C) In the NRe both receptor subtypes are expressed at similar levels. Co-expression of both receptors is rarely observed. Cg, cingulate cortex; PL, prelimbic cortex; IL, infralimbic cortex; mPFC, medial prefrontal cortex; NRe, nucleus reuniens. Graphs show mean ± SEM of mice from all bme points (n=13) since no diurnal differences in OXR1 or OXR2 mRNA expression were evident. **** significant difference between receptors, p<0.0001

**Table 2:** Statistical results for mixed model analysis in medial prefrontal cortex subareas (mPFC). Tukey’s post hoc comparison for percentage of cells containing OXR1 vs. OXR2 mRNA. For the **nucleus reuniens (NRe)**, an unpaired T-test was applied to compare OXR1 vs. OXR2 mRNA expression. Cg, cingulate cortex; PL, prelimbic cortex; IL, infralimbic cortex.

|  |  | Mixed-model analysis/ T-Test | Post hoc OXR1 vs. 2 | Post hoc layer OXR1 | Post hoc layer OXR2 |
| --- | --- | --- | --- | --- | --- |
| **mPFC** | subarea x receptor | F(2,22)=2.096; p=0.1469 | Cg: p<0.0001; PL: p<0.0001; IL: p<0.0001 | Cg vs. PL: p=0.6568; Cg vs. IL: p=0.9961; PL vs. IL: p=0.7872 | Cg vs. PL: p=0.3587; Cg vs. IL: p=0.1118; PL vs. IL: p=0.8878 |
|  | subarea | F(2,22)=0.9206; p=0.4131 |  |  |  |
|  | receptor | F(1,11)=12.85; p=0.0043 |  |  |  |
| **NRe** | receptor | T(12)=2.064; p=0.0613 |  |  |  |

In the mPFC and NRe again only a small fraction of cells expressed mRNA of both receptor subtypes simultaneously (Fig. 2 and S2, table S2).

### 2.3 Comparing diurnal OXR1 mRNA expression levels in the DG hilus

While the number of OXR1(+) cells did not vary significantly over a 24-hour-time course in any of the regions investigated, visual inspection of the sections suggested differences in the number of OXR1 mRNA puncta within single cells of the DG hilus. As an approximation for OXR1(+) expression levels, all OXR1(+) cells were classified as either low (1-5 puncta), moderate (6-10 puncta) or high (> 10 puncta) OXR1 expressing cells. A mixed model analysis for ZT and category of OXR1 content revealed an effect of the OXR1 expression level (Fig. 3; F(2,6)=82.91; p<0.0001) and a significant interaction of OXR1 expression level x ZT (F(6,9)=3.430; p=0.0479; effect of ZT: F(3,9)=3.118; p=0.0809). Tukey’s post hoc comparisons demonstrated a slight reduction of low OXR1 expressing cells from ZT 1 compared to ZT 13 (p=0.0485), suggesting a shift towards moderate to high OXR1 expression levels in the DG hilus during the dark, active period.

**Fig. 3:**
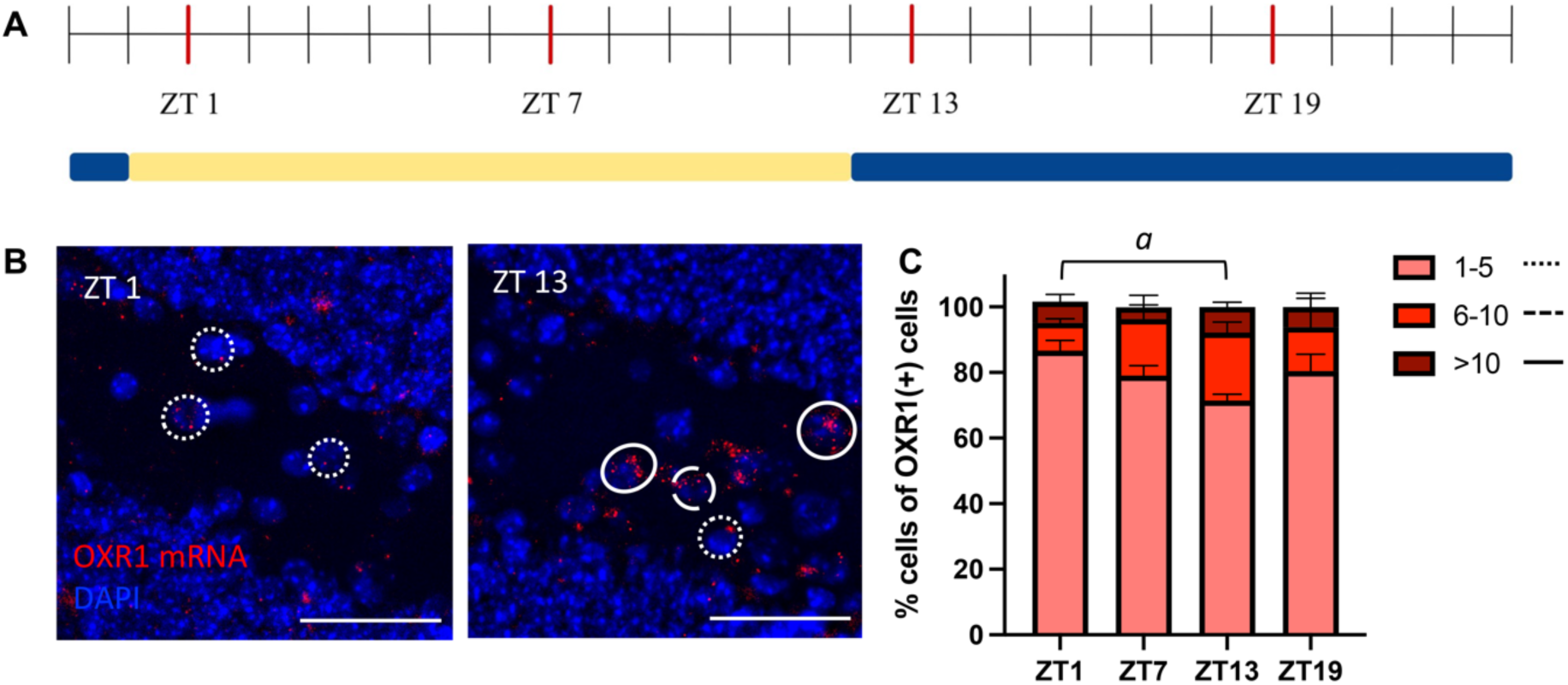
SemiquanLtaLve expression profiles of OXR1 mRNA at different Lmes of perfusion. (A) Time-line indicating the preparation bme points over a 12h light/ 12h dark cycle (Zeitgeber Time, ZT). (B) Example pictures for OXR1 mRNA in the DG hilus are shown at ZT 1 and 13 (Scale bar 50 µm). Cells expressing OXR1 are categorized based on the number of puncta counted in one cells. Examples for low (pointed line), middle (dashed line) and high expressing cells (full line) are marked. (C) Low, middle and high expressing cells expressing OXR1 cells are compared over ZTs. A reduction of low expressing cells towards the beginning of the dark, active phase of the mice are observed. All graphs show mean ± SEM. *a* significant difference between time points for cells expressing 1-5 puncta, p<0.05.

### 2.4 Determining cell types expressing OXR1 mRNA in the DG hilus

The hilus of the dorsal DG is enriched with various types of interneurons that modulate GC activity as well as feedforward and feedback information processing in the DG, determining its output to interconnected brain structures. Based on recent advances in understanding the role of DG interneurons in memory ^27,28^, we selected markers for two different populations of GABAergic interneurons in the hilus to investigate their expression of OXR1 with RNAScope. We observed that only a small fraction of those OXR1(+) cells expressed interneuron markers, without any significant difference in the proportion of those two populations (Fig. 4A; Paired T-Test: T(4)=1.729; p=0.1589).

**Fig. 4:**
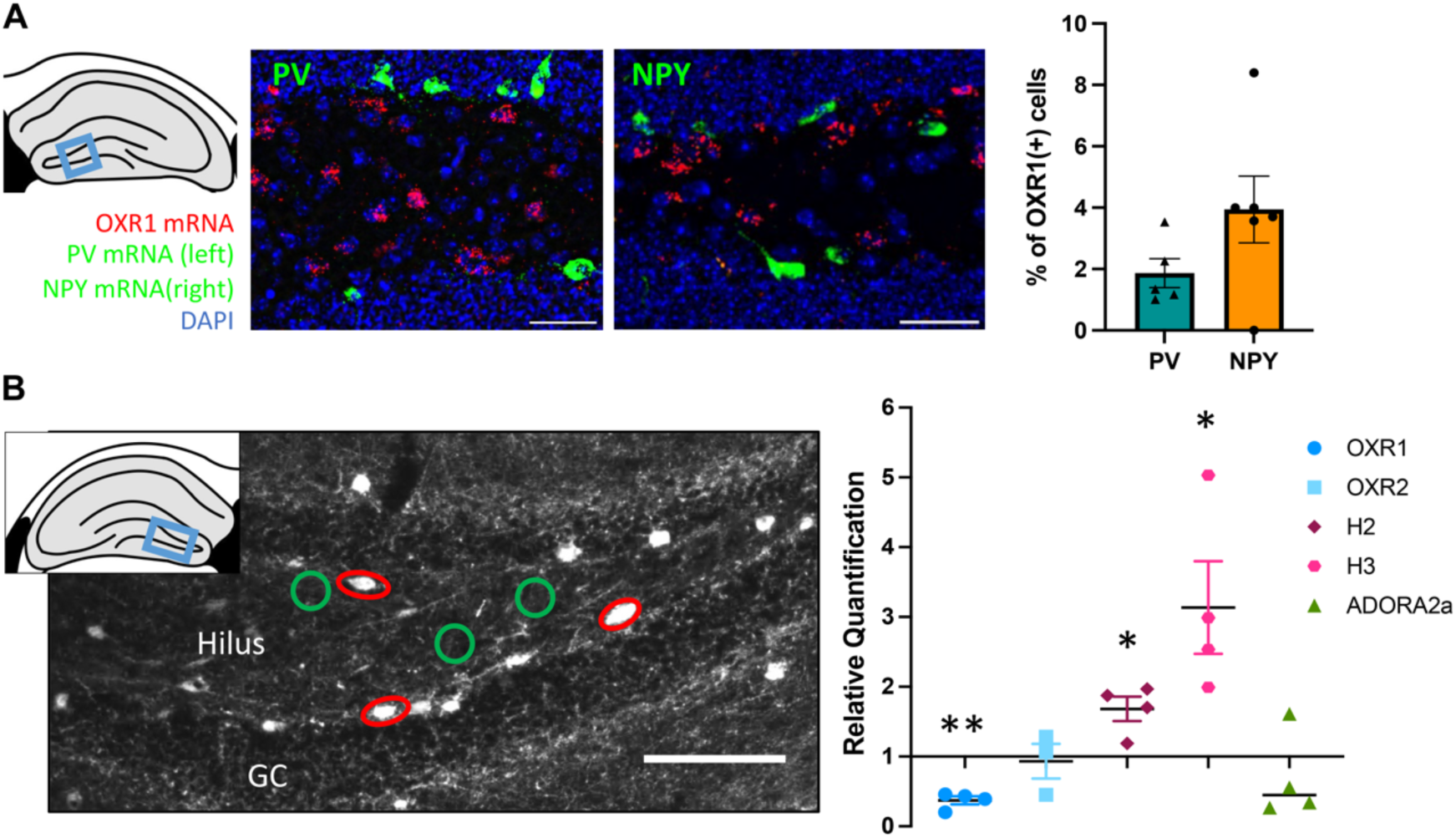
Expression of OXR1 mRNA in DG hilus interneurons. (A) Quantification of the co-expression of parvalbumin mRNA (PV; green, lek) or NPY mRNA (green, right) with OXR1 mRNA (red) demonstrate a low contribution of those interneuron populations to the overall population of OXR1 expressing cells. (B) This was confirmed for NPY-positive interneurons (red circles, lek) using laser microdissection and mRNA expression analysis via quantitative PCR. NPY-positive interneurons express less OXR1 mRNA, but higher mRNA levels of the histamine receptor types 2 and 3 (H2 and H3) compared to NPY-negative cells (green circles; relative quantification (RQ) was set to 1). Comparable mRNA levels for OXR2 and the purinergic receptor ADORA2 were observed in NPY(+) and NPY(-) hilar cells. Scale bar in example images: 50 µm; all graphs mean ± SEM. * significant difference to NPY(-) control tissue, p<0.05; ** p<0.01.

The overall low expression levels of OXR1 mRNA in neuropeptide (NPY) (+) interneurons were confirmed using quantitative polymerase chain reaction (qPCR) of RNA isolated from laser microdissected hilar NPY(+) interneurons marked by GFP in a transgenic mouse line. Here, NPY(+) mRNA levels were normalized to NPY(-) tissue of the hilus (RQ _NPY(-)_ = 1). A One-sample T-Test set against 1 revealed significantly lower OXR1 mRNA in NPY(+) cells (Fig. 4B; T(3)=10.81, p=0.0017), while OXR2 mRNA levels were comparable between NPY(+) and NPY(-) interneurons (T(2))=0.2718, p=0.8112). Since diurnal effects on memory could be mediated by an interplay of other neuromodulators with orexinergic signaling in hippocampus-related brain networks, we also checked for the expression of histamine receptors in hilar NPY(+) interneurons as well as for adenosine receptors. The expression levels of the adenosine receptors were generally low. ADORA2a mRNA did not significantly differ in NPY(+) interneurons (T(3)=0.9782, p=0.4001), while ADORA1 mRNA could not be detected by our qPCR assay. The histaminergic receptor subtypes H2 (T(3)=3.92, p=0.0295) and H3 (T(3)=3.217, p=0.0487) were indeed elevated in NPY(+) interneurons of the hilus compared to NPY(-) hilar tissue, suggesting a susceptibility of this cell type to diurnal neuromodulation beyond orexin.

## 3. Discussion

Using recent advances in fluorescence in situ hybridization techniques, we investigated the expression of the receptor subtypes 1 and 2 for the neuropeptide orexin in sublayers of the dorsal hippocampus, a key structure for spatial and contextual memory, and its interconnected brain areas in C57Bl6J mice. We observed distinct expression patterns for OXR1 and OXR2 mRNA in all regions analyzed with a low portion of cells co-expressing OXR1 and OXR2 (see Fig. 5 for summary).

**Fig. 5:**
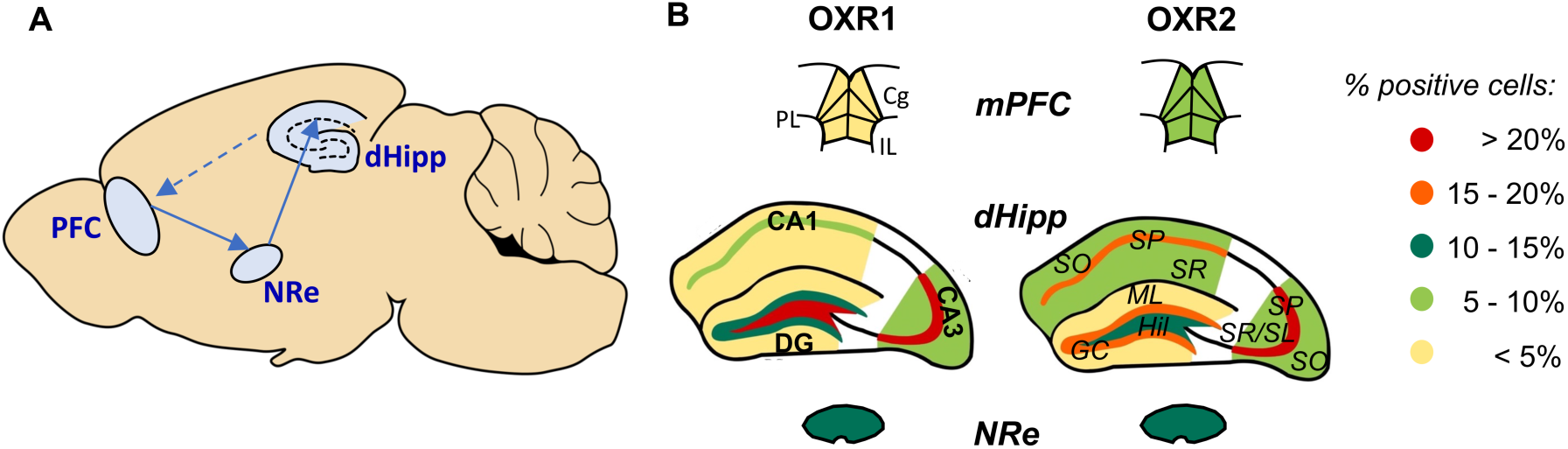
Summary graph for orexin receptor mRNA expression profiles in hippocampal sublayers and selected interconnected areas. (**A**) Scheme representing the connection of the areas analyzed. The ventral part of the hippocampus projects to the mPFC, while major projections from the medial prefrontal cortex (mPFC) to the dorsal hippocampus occur via relay stations such as the midline thalamic nucleus reuniens (NRe). (**B**) The % of cells positive for the orexin receptor (OXR) 1 and OXR2 mRNAs are distinctly distributed in the sublayers of the dorsal hippocampus and in subareas of the hippocampus, is indicated by the color code. Cg, cingulate cortex; DG, dentate gyrus; GC, granular cell layer; IL, infralimbic cortex; ML, molecular cell layer; CA, cornu ammonis; PL, prelimbic cortex; SO, stratum oriens; SP, stratum pyramidale; SR, stratum radiatum; SL, stratum lucidum.

The hippocampal expression profiles of OXR1 and OXR2 mRNA in mice are comparable to data available in rats ^22,23^, describing low to moderate expression levels for OXR1 mRNA in the rats dorsal DG and CA1 area and high to moderate OXR2 mRNA levels in the CA3. On the protein level, a strong expression of OXR2 in CA3 pyramidal cell somata and moderate OXR1 levels in in DG granule cells and in the hilus of rats are confirmed by immunohistochemistry ^24,25^. In transgenic mice, OXR1-GFP-positive somata are observed especially along the pyramidal cell layer of CA1 and CA3 and within the dentate gyrus ^29^. A very recent study using another high sensitivity in situ hybridization technique for mapping OXR1 and OXR2 mRNA throughout the mouse brain confirms overall the expression patterns obtained by us in the hippocampal sublayers, indicating moderate OXR1 levels in the granule cell layer, in the hilus of the dorsal DG and in the *stratum pyramidale* of the CA3. As in our data, the expression of OXR2 mRNA is highest in the CA3 subarea ^26^.

In the mPFC, we observed a significant higher expression of OXR2 compared to OXR1 mRNA throughout the Cg, PL and IL subregions. While these regions are not analyzed in detail in the recent study by Tsuneako & Funato, their representative images suggest a similar pattern ^26^, which is futher confirmed in transgenic OXR1-GFP mice ^29^. In rat brains, OXR2 mRNA is primarily described to be present in the cingulate and other neocortical areas, specifically in layer IV and VI ^22,23^, but such differences may stem from dissimilar approaches for defining areas of interest (e.g. assessment of the whole frontal cortex vs. mPFC subareas).

Within the NRe, we detected low OXR1 and OXR2 mRNA levels, which matches observations made in the rat brain ^23^. While Tsuneoka & Funato describe no expression of OXR1 mRNA in the NRe beyond background levels ^26^. Nevertheless, there are few cells positive for OXR1 visible in their representative images depending on the plane analyzed. Our analysis approach of counting the fraction of positive cells for OXR1 and OXR2 mRNA in a larger cohort of mice may provide a higher sensitivity under low expression conditions. Cells expressing both receptor subtypes simultaneously were rarely observed. This is in agreement with the results of the brain wide analysis demonstrating a co-expression of OXR1 and OXR2 mRNA only in the paraventricular thalamus, the ventromedial nucleus of the hypothalamus and the dorsal raphe nucleus of the brain stem ^26^.

High sensitivity in situ approaches like RNAScope or the branched in situ hybridization chain reaction (bHCR) technique utilized by Tsuneoka & Funato ^26^ enable the detection of few mRNA molecules and the number of puncta detected in a cell corresponds to a certain extend to the amount of mRNA within a cell. Using both techniques, the number of puncta usually detected within a single cell is overall low (1-5 puncta in most cells; Fig. 1+2, ^26^). The number of OXR1 or OXR2 mRNA positive cells did not differ between preparation time points over a 12h dark/ 12h light cycle. To assess whether differences occur rather in the amount of OXR1 or OXR2 mRNA produced within a cell, a high-resolution analysis estimating expression intensities was performed for OXR1 mRNA in the dorsal DG hilus. This region showed an overall moderate number of OXR1 positive cells. We observed a slight shift towards cells expressing more OXR1 mRNA puncta at the beginning of the dark, active phase (ZT 13) compared to the beginning of the light, inactive phase (ZT 1), aligned to the activity peak of orexinergic neurons in the lateral hypothalamus ^8^. Using qPCR, a peak in hippocampal OXR1 and OXR2 mRNA levels has been observed in the middle of the dark phase, thereby correlating positively with the expression pattern for circadian clock genes such as bmal1 ^18^. Similar associations are observed for OXR2 in the hypothalamus and cortex ^30^. A qPCR approach would allow to detect diurnal quantitative shifts in mRNA levels more precisely, but with the downside of a low spatial resolution and a lack of cell-type specificity.

The hilar region of the DG contains diverse populations of interneurons that control hippocampal circuit activity ^31^. As the two major populations of GABAergic interneurons, parvalbumin (PV) cells provide a proximal inhibition of DG granule cells and drive feedforward inhibition within the DG-CA3 circuit, while hilar perforant path associated (HIPP) cells, marked by their expression of somatostatin (SST) and NPY, provide a distal inhibition of granule cells ^28,32^. Both cell types control the precision of contextual fear memories, either under remote condition ^33^ or by confining DG granule cell ensembles recruited to a contextual memory engram ^27,34^. Acute sleep deprivation disturbs contextual fear memory consolidation by over-activating HIPP cells ^34^. Thus, a direct modulation of HIPP cells by orexin might be possible, but we could not observe a particular enrichment of OXR1 mRNA in this cell type, using NPY as a marker. Similarly, only a minor fraction of hilar cell containing OXR1 mRNA co-expressed PV mRNA. We then used qPCR in laser-microdissected NPY(+) HIPP cells to analyze OXR1 and OXR2 mRNA expression levels. With this technique we confirmed that HIPP cells express significantly less OXR1 than NPY(-) hilar control tissue, suggesting prevalent OXR1 expression in other cell populations of the hilus. Nevertheless, HIPP cells showed an enriched expression of the histamine receptors H2 and H3. Since histamine release in the hypothalamus is triggered by orexin ^36^, HIPP cells may be indirectly affected by orexin via a histaminergic stimulation. In addition, NPY(+) cells expressed OXR2 and the adenosine receptor type 2a (ADORA2a) at similar levels compared to NPY(-) control tissue in this area, allowing for a modulation of local circuits involving HIPP cells via these receptors although the overall expression of OXR2 mRNA in the hilus is low. Like HIPP cells, adenosine type 2a receptors control contextual fear memory specificity ^37^ and further studies are needed to investigate the exact mechanisms and local circuits of such a modulation.

Overall, both orexin receptor subtypes were expressed in all subfields of the dorsal hippocampus and its interconnected areas, mPFC and NRe, with different densities (see Fig. 5 for summary). So far, pharmacological studies have helped to address the functional implications of such expression patterns. In the hippocampus, OXR1 activation supports the induction of long-term potentiation (LTP) at DG perforant path-granule cell synapses ^38^ and at CA1 Schaffer collaterals ^39^, but re-potentiation in the CA1 under low frequency stimulation requires both orexin receptors ^40^. Activation of OXR1 in the dorsal CA1 and the DG is required for the acquisition and consolidation of spatial memory in a water maze task, while effects on its retrieval are restricted to the CA1 but not the DG region ^15,16^. Accordingly, a local shRNA-mediated knock down of OXR1 in the CA1 region impairs spatial learning in the water maze in Nile grass rats ^41^. Cued and contextual fear memory acquisition are impaired after systemic administration of OXR1 and OXR2 antagonists ^14^, while contextual fear memory retrieval is only reduced by OXR1 but not OXR2 antagonist ^42^. Contextual aversive memory is dysregulated in post-traumatic stress disorder (PTSD). In a PTSD rat models, an increased expression of hippocampal OXR1 mRNA is observed, but these animals had no long-term dysregulation of contextual fear memory, suggesting a contribution of this receptor subtype to resilience to traumatic stress and the maintenance of precise fear memories ^17^. In animal models of Alzheimer’s disease, a disorder characterized by early hippocampus-dependent memory impairments, hippocampal OXR1 and OXR2 mRNA levels are reduced especially during the dark, inactive phase ^18^. However, the treatment of Alzheimer’s disease mouse models with OXR1 and OXR2 antagonists elicits contradictory effects on memory performances so far ^43,44^.

Within the mPFC, OXR1 and OXR2 both functionally contribute to emotional states, for example by enhancing anhedonia in a chronic unpredictable stress model of depression in mice ^45^, and by promoting the tolerance to threatening stimuli via orexinergic signaling in the PL subregion ^46^. In the Cg subarea, OXR1 is engaged in decision-making processes in cost-benefit-dependent working memory tasks ^47^. OXR1 in the mPFC modulates reward-related behavior ^48^, while OXR2 can potentially modualte attention and arousal by fine-tuning cortical oscillations, as shown for thalamocortical networks involving neocortical layer 6b neurons of the sensory cortex ^20^. In the thalamus, the paraventricular nucleus is the main target of orexinergic projections and contributes to reward-associated behaviors via OXR2 signaling ^49^. Whether orexinergic signaling in the NRe would contribute to learning and memory warrants further investigation.

In summary we applied RNScope as a commercially available, advanced tool for high-sensitivity fluorescence in situ hybridization labeling to investigate the expression of the receptor subtypes 1 and 2 for the neuropeptide orexin in C57Bl6J mice. We focused on sublayers of the dorsal hippocampus and the PFC and NRe as key network structures of working and spatial as well as contextual memory. We observed differential, layer-specific expression of OXR1 and OXR2 mRNA in the dorsal DG, CA3 and CA1 region as well as in the mPFC with a relative increase in the density of OXR1 compared to OXR2 mRNA particularly in the hilus of the DG. Here, a minor shift of OXR1 mRNA towards higher expression levels at the beginning of the active phase was observed. While functional studies suggest a primary role of OXR1 in hippocampus-dependent spatial and contextual memory, the exact contribution of orexin signaling via OXR1 to local circuit activity within the DG-CA3-CA1 subfields remains unclear. We began to address this question by applying a cell-type specific RNAScope and qPCR analysis of OXR1 mRNA expression for two major populations of GABAergic interneurons shaping contextual memory engrams, but found no enrichment. Deciphering the detailed orexinergic modulation of local circuits in the hippocampus and its interconnections with the mPFC, both directly and via the NRe, remains an exciting subject of future studies. Orexin may indeed be a “key to a healthy life” ^1^, once the neurobiological basis of an orexinergic modulation of entities like memory and emotion regulation can be clarified.

## 4. Methods

### 4.1 Animals

Male C57BL/6 mice and NPY-GFP mice (B6.FVB-Tg(Npy-hrGFP)1Lowl/J) were bred and raised at the central animal facility of the medical faculty of the Otto-von-Guericke University Magdeburg. They were kept in groups of 2-3 animals under a 12:12 inverse light-dark regime with lights off at 7:00 am and lights on at 7:00 pm and with water and food *ad libitum*. Animal housing and all experiments were performed in accordance with the European and German regulations, approved by the Landesverwaltungsamt Sachsonia-Anhalt (Permission Nr. 42502-2-1717) and following the ARRIVE guidelines. Only young adult (10-16 weeks) male mice were used for the experiments.

### 4.2 Fluorescence in situ hybridizaBon (RNAScope)

#### Tissue Collection & Preparation

The brains of 13 C57BL/6J mice were prepared at four time points spaced 6 hours apart over the course of 24 hours (ZT 0= lights on). The first time point (ZT 1, n=4) was sampled one hour after the onset of light (start of the inactive phase) while ZT 13 (n=3, as for ZT8, ZT, 19) occurred one hour after the offset of light (start of the active phase). Naïve NPY-GFP mice (n=4) and an additional set of C57Bl6J mice (n=6) for the co-labelling experiment of OXR1 and interneuron markers were prepared at ZT 19. Mice were transcardially perfused with PBS followed by 4% PFA under deep anaesthesia, the brains were then removed and postfixed overnight in 4% PFA at 4°C, followed by immersion in 20% sucrose solution. All brains were snap frozen in 2-methyl butane cooled in dry ice and stored at −80°C until cryosectioning commenced under RNAse free conditions. Coronal sections were cut (10µm) at the level of the mPFC (Bregma 1.94 to 1.34 mm), the NRe (Bregma −0.58 to −1.06 mm) and the dorsal hippocampus (Bregma −1.46 to −2.18 mm; all according to ^50^). Brain sections were directly transferred to pre-baked Super Frost Plus slides (Thermo Scientific,Braunschweig, Germany). Coronal sections of NPY-GFP mice were mounted on membrane covered slides to allow for laser capture microdissection (Membrane Slide 1.0 PEN, Zeiss, Wetzlar, Germany). The slides were dried and stored at −80°C until further use.

#### Fluorescence in situ hybridization (RNAScope)

Fluorescence in situ hybridization was performed according to the manufacturer’s instructions for RNAscope Multiplex Fluorescent Assay Version 1 (Cat. No 320850) and Version 2 (Cat. No 323100) for fixed frozen brain tissue (Advanced Cell Diagnostics (ACD), Newark, CA, USA;^51^). All slides were pre-treated by baking (10-30min, 60°C), post-fixation (15min, 4°C, 4% PFA), dehydration (50% EtOH, 70% EtOH, 100% EtOH), hydrogen peroxide incubation, target retrieval (5/15min, 98°C) and protease III treatment (30min, humid chamber, 40°C).

For detecting ***OXR1 and OXR2 mRNA co-labelling,*** probe hybridization was performed by incubating the slides with the C3 probes (OXR2: Probe-Mm-Hcrtr2-O1-C3, #581631-C3, ACD) diluted in the C1 probe (1:50; OXR1: Probe-Mm-Hcrtr1, #466631, ACD) for 2h at 40°C in humid conditions, followed by hybridization of the probe with Amplifier 1 and subsequent binding of Amplifier 2-4. A fluorescent dye was connected to Amplifier 4 (RNAscope assay version 1: Amp 4 Alt C-Fl; OXR1-C1 probe linked to Atto 550; OXR2-C3 probe linked to Alexa 488).

For detecting ***OXR1 mRNA co-expression in interneuron subpopulations*** of the DG hilus, slices of the dorsal hippocampus were similarly treated with probes for OXR1 (C1; Probe-Mm-Hcrtr1, #466631, ACD) and NPY (C2; Probe-Mm-Npy-C2, #313321, ACD) and incubated with the amplifiers, linked to the fluorescent dye in the last step (Amp 4 Alt B-Fl (assay version 1; OXR1-Atto 550 + NPY-Alexa 488). A separate set of slides was incubated with a probe set for OXR1 and parvalbumin (C3; Probe-Mm-Pvalb, #421931, ACD). Here, due to discontinuation of the RNAScope Multiplex Fluorescent Assay version 1, version 2 was used, where the fluorescent signal was developed for each channel separately using Tyramide Signal Amplification (TSA) technology (assay version 2: OXR1-TSA-570 + PV-TSA-650).

All experiments included one slide each for positive and negative control probe incubation (RNAscope 3-plex Positive Control Probe #320881; Negative Control Probe, #320881; ACD). Two to three slides per animal were labelled and all slides were incubated with DAPI provided by the kit (30sec, #323108, ACD) for nulcei staining before embedding and cover slipping with ProLong Gold Antifade Mountant (P36930, Invitrogen, Eugene, USA).

#### Microscopy & image analysis

Images were acquired with an epifluorescence microscope (Leica THUNDER DMi8, Wetzlar, Germany) with Instant Computational Clearing (ICC, refractive index: 1.4700, 40x objective) and analysed using the open-source software QuPath ^52^. ROIs were placed manually lining out the sublayers of the dorsal hippocampus (hilus, granule cell layer and molecular layer of the dentate gyrus; *stratum radiatum/ lucidum*, *pyramidale* and *oriens* of CA3; *stratum radiatum, pyramidale* and *oriens* of CA1; ^50^). For the mPFC and the NRe ROIs were positioned based on the size and location of the brain region in the given bregma plane. First, the total cell number of a selected region was determined in the DAPI channel by applying the Cell Detection Tool (settings: Requested pixel size (0.5µm), Background radius (8µm), Median filter radius (0µm), Sigma (1.5µm), Minimum Area (20 µm^2^), Maximum Area (400µm^2^), complemented with the soma extension option (5µm) and corrected manually if needed. The Intensity Parameter Threshold which was adjusted to optimize the cell recognition (ranging 10 – 30). OXR1 and OXR2 single-positive and double-positive cells were counted manually in a given region and sublayer. The percentage of cells expressing either one of the target mRNAs or both was calculated for each region and per animal covering 2-3 slides with both hemispheres each. To assess the OXR1 mRNA expression level of the hilus in higher detail, the exact number of puncta was counted in a representative subset of hilar cells provided by the Cell Detection Tool.

### 4.3 QuanBtaBve PCR in hilar NPY-posiBve interneurons

#### Quantitative PCR in Laser Capture Microdissection (LCM), RNA isolation and cDNA synthesis

As described previously ^27^, PFA-fixed frozen slices (14 µm) of dorsal hippocampus of NPY-GFP mice were mounted on pLL-coated RNAse-free membrane slides and manually marked under the fluorescence mode of PALM MicroBeam system (Carl Zeiss, Jena, Germany) for subsequent laser microdissection of single NPY(+) cells from the DG hilus (about 400 cells per sample covering 91000 to 93000 µm2). NPY(-) cells were collected by dissecting non-fluorescent areas of the same size within the hilus afterwards. Sample lysis and the isolation of total RNA isolation was performed with the RNeasy® FFPE kit (Qiagen, Hilden, Germany) according to the manufacturer’s instructions. First, strand cDNA was synthetized using the TaKaRa PrimeScipt RT Reagent Kit (Cat# RR037A, Takara Bio Inc., Kusatsu, Japan) using 20 µM Random decamer first strand primers according to the manufactorer’s instructions.

#### Quantitative PCR & data analysis

A duplex real-time PCR (ABI Prism Step One real-time PCR apparatus, Thermo Fisher Scientific, Waltham, MA, USA) was performed in triplicates of 1:10 diluted cDNA samples with predesigned FAM-labeled TaqMan® assays (Applied biosystems by Thermo Fisher Scientific, Waltham, MA, USA) for OXR1 (Assay ID: Mm01185776_m1), OXR2 (Mm0179312_m1), histamine receptor type 2 (H2; Mm00434009_s1), histamine receptor type 3 (H3; Mm00446706_m1), adenosine receptor 2a (ADORA2a; Mm00802075_m1) and a VIC-labelled assay for the housekeeping gene glycerinaldehyd-3-phosphat-dehydrogenase (GAPDH; endogenous control; Life Technologies), running 50 cycles of 15 s at 95°C and 1 min at 60°C, preceded by a 2 min 50°C decontamination step with Uracil-N-glycosidase and initial denaturation at 95°C for 10 min. Relative quantification of gene expression was achieved using the ddCT method by first normalizing the mean cycle threshold (CT) to the internal control GAPDH for each sample (dCT; dCT = CT_target_ − CT_GAPDH_). All samples were then normalized to the mean dCT value for the respective NPY(-) samples by calculating ddCT = dCT_sample_ – mean dCT_NPY(-)_. Lastly, the relative quantification (RQ) was calculated using the following formula: RQ = 2^-ddCT^, yielding a normalized expression of NPY(-) cells =1 for the respective target.

### 4.4 StaBsBcal Analysis

Statistical analysis was performed using GraphPad Prism (GraphPad Software Inc., Version 9). Normal distribution of values was tested with the Shapiro-Wilk test. For analyzing OXR1 and OXR2 mRNA expression profiles across hippocampal sublayers, within mPFC subareas and in the NRe, a mixed model analysis with Geisser-Greenhouse corrections was used to account for unequal variability of differences. First, main effects of hippocampal layer/ subarea, ZT and their interaction were investigated separately for the population of OXR1, OXR2 and double-positive cells. After a lack of significant ZT effects, ZT groups were combined and the factor “receptor” (OXR1 vs. OXR2) was introduced to the mixed-model analysis for their main effects and interactions with the sublayers/ subareas. Posthoc analysis was performed with Tukey’s multiple comparison test. Similarly, for the semiquantitative analysis of OXR1 expression categories in the DG hilus, a mixed-effect analysis followed by Tukey’s posthoc comparison was performed for the factors ZT and expression category. For the relative quantification of target gene expression in NPY(+) cells, a one-sample t-test against 1, as the set RQ value for NPY(-) cells, was performed.

## Data availability statement

The datasets generated during and/or analyzed during the current study are available from the corresponding author on reasonable request.

## Acknowledgement

We thank Romina Wolter and Annika Lenuweit for excellent technical assistance. This work was supported by a grant from the German Research Foundation (Project-ID 425899996 – CRC 1436) to Anne Albrecht and from the Center for Behavioural Brain Sciences Magdeburg - CBBS funded by the European funds for regional development (EFRE, Funding Nr. ZS/2016/04/78113).

## Author contributions

AA, GMK and LMCB conceptualized the work; AA provided resources and acquired the funding; GMK and LMCB acquired and analyzed the data with support from AA; AA wrote the main manuscript text, GMK prepared figures 1-5 with support from LMCB for Fig. 4. All authors reviewed and edited the manuscript.

## Competing interest

The authors declare no competing interest

## Notes

### Competing Interest Statement

The authors have declared no competing interest.

### Summary of Updates

In the previous version of the manuscript we adhered to the more general, historical definition of a circadian phase, considering a phase spanning an approximately 24h cycle, given here by light as a Zeitgeber in our a 12h dark/12light rearing conditions. To adhere to the stricter definition by Viterana et al. (2001, PMID: 11584554), we will refer now in our observations to the diurnal rhythm, since we did not test whether any potential time-of-day dependent expression modulation of OXR1 and OXR2 mRNA was endogenous and independent of a Zeitgeber such as external light. This was indeed not the aim of the current study. Rather, we focused on investigating high resolution spatial expression profiles of orexin receptors in memory-relevant areas. But since orexin neurons show activity differences dependent on the wake/ sleep-cycle and the time of the day, we considered such potential effects as well when assessing the expression profiles. To account for the 12h dark/ 12h light rhythm used in our study and different preparation time points therein, we now carefully adapted the manuscript, including the supplementary material, and referred to diurnal phases when appropriate. We furthermore shortened the discussion and improved the contrast in the microscopic example pictures.

