## Supplementary Material for "Spatial mRNA expression patterns of orexin receptors in the dorsal hippocampus"

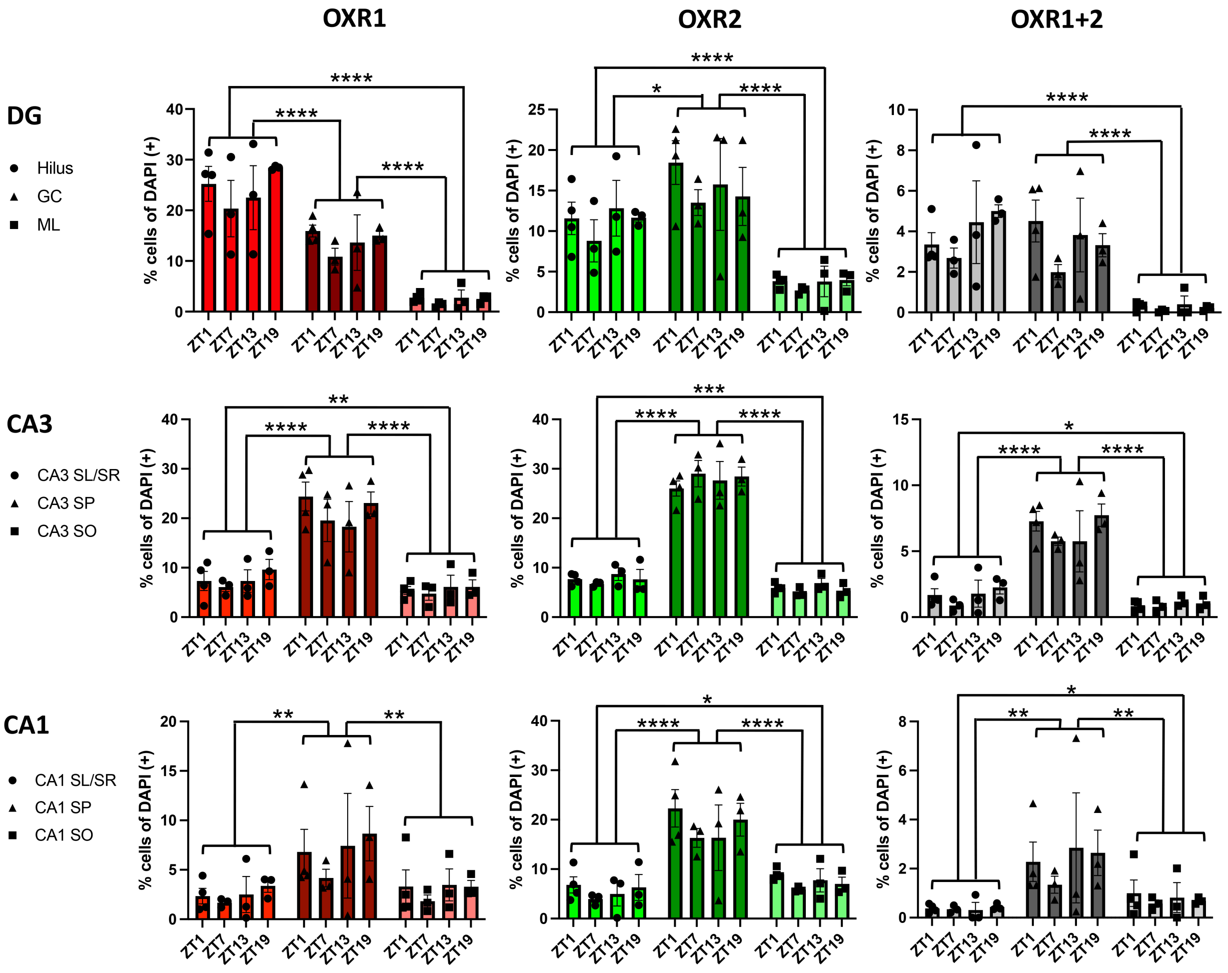


**Supplementary Figure S1: Diurnal orexin receptor mRNA expression profiles in hippocampal sublayers.** The expression levels of OXR1 and OXR2 show layer-specific patterns in the dorsal hippocampus, but no significant effects of the time point of preparation (ZT 0 = lights on in a 12h/ 12h light dark circle). The portion of cells co-expressing OXR1 and OXR2 is low in all areas. DG, dentate gyrus; GC, granular cell layer; ML, molecular cell layer; CA, cornu ammonis; SO, stratum oriens; SP, stratum pyramidale; SR, stratum radiatum; SL, stratum lucidum. Graphs show mean ± SEM with n=3-4 per time point. ** significant difference between layers, ** $p<0.01$; *** $p<0.001$; **** $p<0.0001$.

**Supplementary Table S1: Statistical results for mixed model analysis of diurnal effects in hippocampal sublayers.** Zeitgeber time (ZT) indicates 4 different time points over a 12h lights / 12 h dark period (ZT0 = light on). Differences in OXR1 mRNA, OXR2 mRNA and co-expression of both receptor mRNAs were analyzed separately.
DG, dentate gyrus; GC, granular cell layer; ML, molecular cell layer; CA, cornu ammonis; SO, stratum oriens; SP, stratum pyramidale; SR, stratum radiatum; SL, stratum lucidum.

| | | **OXR1** | **OXR2** | **OXR1+2** |
|---|---|---|---|---|
| **DG** | ZT x layer | F(6,18)=0.6168; p=0.7143 | F(6,18)=0.4536; p=0.8331 | F(6,18)=1.613; p=0.2007 |
| | ZT | F(3,9)=0.6046; p=0.6284 | F(3,9)=0.4156; p=0.7460 | F(3,9)=0.6764; p=0.5880 |
| | layer | F(1.258,11.32)=106.8; p<0.0001 | F(1.568,14.11)=40.80; p<0.0001 | F(1.734,15.60)=43.70; p<0.0001 |
| **CA3** | ZT x layer | F(6,18)=1.259; p=0.3243 | F(6, 18)=0.6775; p=0.6697 | F(6,18)=0.7826; p=0.5945 |
| | ZT | F(3, 9)=0.3564; p=0.7859 | F(3,9)=0.1743; p=0.9111 | F(3,9)=0.6357; p=0.6106 |
| | layer | F(1.230,11.07)=128.9; p<0.0001 | F(1.209,10.89)=349.0; p<0.0001 | F(1.245,11.20)=96.43; p<0.0001 |
| **CA1** | ZT x layer | F(6,18)=0.3204; p=0.9177 | F(6,18)=0.4327; p=0.8475 | F(6,18)=0.3780; p=0.8834 |
| | ZT | F(3,9)=0.3541; p=0.7875 | F(3,9)=0.5879; p=0.6381 | F(3,9)=0.1811; p=0.9066 |
| | layer | F(1.184,10.65)=13.07; p=0.0033 | F(1.199,10.79)=57.37; p<0.0001 | F(1.099,9.895)=11.70; p=0.0058 |

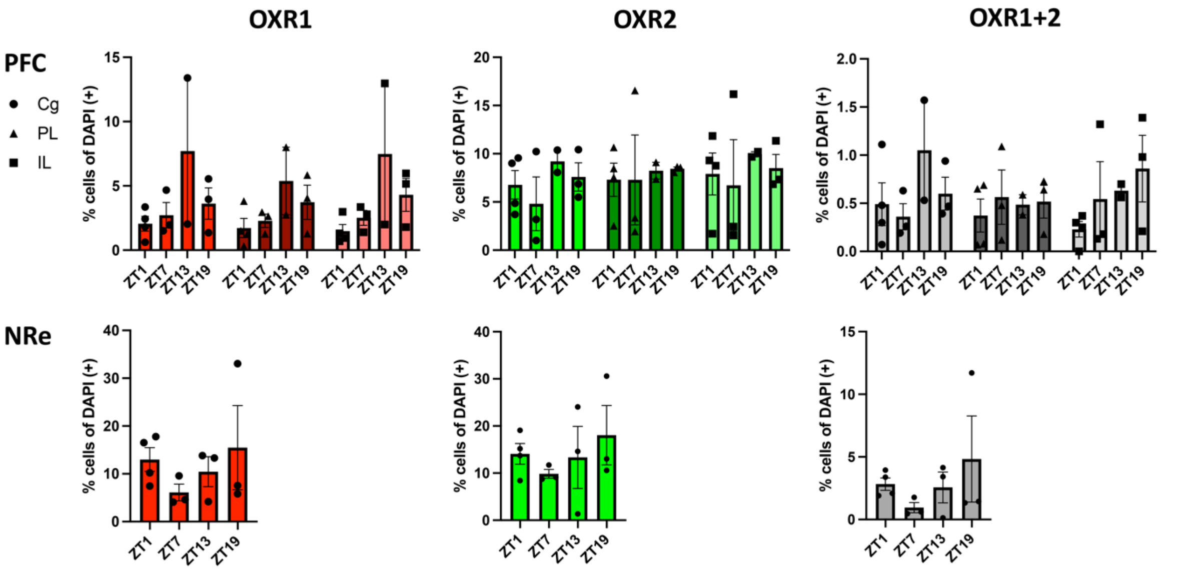


**Supplementary Figure S2: Diurnal orexin receptor mRNA expression profiles in the medial prefrontal cortex subregions and nucleus reuniens relay station to the hippocampus.** All subregions of the mPFC show similar levels of OXR2 and OXR1 mRNA expression densities. Within the mPFC sublayer and in the NRe no significant effects of the time point of preparation were evident (ZT 0 = lights on in a 12h/ 12h light dark circle). Co-expression of both receptors is rarely observed. Cg, cingulate cortex; PL, prelimbic cortex; IL, infralimbic cortex; mPFC, medial prefrontal cortex; NRe, nucleus reuniens. Graphs show mean ± SEM with n=3-4 per time point.

**Supplementary Table S2: Statistical results for mixed model analysis of diurnal effects in medial prefrontal cortex (mPFC) subareas and in the nucleus reuniens (NRe).** Zeitgeber time (ZT) indicates 4 different time points over a 12h lights / 12 h dark period (ZT0 = light on). Differences in OXR1 mRNA, OXR2 mRNA and co-expression mRNA for both receptors were analyzed separately. For the nucleus reuniens only main effects of ZT were analyzed for OXR1, OXR2 and their combined mRNA expression.
Cg, cingulate cortex; PL, prelimbic cortex; IL, infralimbic cortex.

| | | **OXR1** | **OXR2** | **OXR1+2** |
|---|---|---|---|---|
| **PFC** | ZT x subarea | F(6,16)=0.7280; p=0.6338 | F(6,16)=0.4301; p=0.8482 | F(6,16)=1.601; p=0.2107 |
| | ZT | F(3,8)=1.790; p=0.2268 | F(3,8)=0.2203; p=0.8796 | F(3,8)=0.6291; p=0.6164 |
| | subarea | F(1.559,12.47)=1.361; p=0.2844 | F(1.616,12.93)=1.268; p=0.3056 | F(1.272,10.18)=0.7913; p=0.4245 |
| **NRe** | ZT | F(3,9)=0.7156; p=0.5671 | F(3,9)=0.5447; p=0.6639 | F(3,9)=0.8134; p=0.5180 |
